# Design and implementation of an intelligent framework for supporting evidence-based treatment recommendations in precision oncology

**DOI:** 10.1101/2020.11.15.383448

**Authors:** Frank P.Y. Lin

## Abstract

**BACKGROUND:** The advances in genome sequencing technologies have provided new opportunities for delivering targeted therapy to patients with advanced cancer. However, these high-throughput assays have also created a multitude of challenges for oncologists in treatment selection, demanding a new approach to support decision-making in clinics.

**METHODS:** To address this unmet need, this paper describes the design of a symbolic reasoning framework using the method of hierarchical task analysis. Based on this framework, an evidence-based treatment recommendation system was implemented for supporting decision-making based on a patient’s clinicopathologic and biomarker profiles.

**RESULTS:** This intelligent framework captures a six-step sequential decision process: (1) concept expansion by ontology matching, (2) evidence matching, (3) evidence grading and value-based prioritisation, (4) clinical hypothesis generation, (5) recommendation ranking, and (6) recommendation filtering. The importance of balancing evidence-based and hypothesis-driven treatment recommendations is also highlighted. Of note, tracking history of inference has emerged to be a critical step to allow rational prioritisation of recommendations. The concept of inference tracking also enables the derivation of a novel measure — level of matching — that helps to convey whether a treatment recommendation is drawn from incomplete knowledge during the reasoning process.

**CONCLUSIONS:** This framework systematically encapsulates oncologist’s treatment decisionmaking process. Further evaluations in prospective clinical studies are warranted to demonstrate how this computational pipeline can be integrated into oncology practice to improve outcomes.

## 1 Introduction

Decision-making in oncology is increasingly reliant on nuanced patient characteristics and sophisticated biomarkers. At the point of care, there is a fundamental need for both precision and efficiency. In the era of rapid therapeutic advances, the ever-increasing information burden has limited the practicality of conducting rigorous review of evidence at cursory physician-patient encounters. Excessive cognitive load is also often placed on physicians, leading to application of various heuristics to improve efficiency at the expense of optimal decision-making in the vast therapeutics landscape.

While consensus-based guidelines have been established to mitigate physician’s information needs, there is often lack of standardised protocol to guide drug selection in the treatment-refractory settings. Increasingly, oncologists have access to massively parallel sequencing assays to gain a glimpse of druggable targets in a patient’s tumour [1]. In large-scale precision oncology programs, patients matched to therapies selected by next-generation sequencing (NGS) have comparatively better tumour response to treatments; clinical benefits have also been shown in some reports [2,3]. However, the overall magnitude-of-benefit of using large-scale tumour genotyping to find a druggable biology remains small [4,5], with expert groups calling for its use only in limited clinical scenarios [6]. In addition to the issue of patient selection, there are operational challenges to bring NGS-based assays to the bedside, including but not limited to sampling issues, computational limitations, variant interpretation, biomarker complexity, as well as various financial, and administrative barriers [1,7–9].

One major hurdle is that large NGS panels often find more potentially “actionable” molecular targets than our knowledge about the drugs. Many clinical dilemmas are also created when making drug recommendations based on inadequate efficacy and safety data. Practically, timely and accurate appraisal of evidence often forms a significant barrier to realise NGS technology to its full potential. When making treatment decisions, skilled oncologists often need to navigate through numerous “evidence-gaps”. Not surprisingly, divergent opinions are often seen between expert groups [10], highlighting the need for a standardised, more systematic solution to decision-making.

Algorithmic decision support holds the promise to reduce physician’s cognitive load, and its use has been suggested to improve the quality of shared decision process in tumour boards [11–14]. Such digital platforms, now often incorporated into “virtual” molecular tumour boards, are increasingly used to translate results of genomic profiling to help therapy selection [15,17]. Herein, a framework for developing an intelligent agent, using a symbolic artificial intelligence (AI) method for decision support, is proposed to address this critical information need in oncologist’s treatment recommendation process. From a technical perspective, while methods of symbolic reasoning are well-established, there is no consensus on how clinical expertise in precision oncology can be readily shared using digital platforms. Thus, the structure of this intelligent framework also serves to standardise the task of supporting evidence-based, molecularly informed recommendations.

## 2 Methods

### 2.1 Design of the intelligent framework

Hierarchical task analysis was used to produce a clinical workflow model [16], based on the reasoning process of a medical oncologist (FL) in making therapy recommendations for patients with advanced cancer. To build a clinical recommendation framework, the top-level hierarchy derived from the task decomposition process was assigned as the core steps of this framework. A list of clinical premises, oncologist’s heuristics, and output requirements associated with a functional recommendation system were also formulated by the author.

### 2.2 Implementation and validation of the framework

To demonstrate the utility of this framework, a treatment recommendation system was implemented using the framework structure and the requirements elicited above. An expert system shell was implemented in Perl programming language (version 5) to handle symbolic inference based on production rules. Case studies were reported to illustrate how recommendations of targeted therapies can be prioritised based on cancer types and results of and molecular alterations of the tumour.

## 3 Results

### 3.1 An *in silico* framework for oncology treatment recommendations

#### 3.1.1 Design of an intelligent agent and task environment

##### The intelligent agent (IA)

The requirements of an Intelligent Agent (IA) form the basis of an implementable Treatment Recommendation System (TRS). The IA should recapitulate oncologists’ decision process to produce rational recommendations based on a patient’s clinicopathologic characteristics. Closely aligned to oncologists’ decision-making process, several heuristics are readily recognised when formulating a treatment plan for patients with metastatic cancer after standard-of-care options are exhausted, which should be modelled by the TRS (Box 1).

###### Box 1 List of premises in making treatment decisions for patients with refractory cancer.

- Therapy (including therapy combinations) recommendations should be rationally prioritised based on the preceived clinical value of a therapy, which integrates aspects of efficacy, safety, availability, and biological activities.
- The strength-of-recommendation and clinical value of a therapy option can be approximated by summarising the regulatory approval status, results from rationally conducted clinical trials, recommendation by consensus guidelines, in place of direct assessment of efficacy, safety, availability, and biological activities of a drug.
- A therapy (or therapy combination) in development from the same drug class of another agent with a higher LOE, is more likely to achieve therapeutic significance when compared to an agent from a less established drug class.
- In the absence of direct evidence to report, level-of-evidence (LOE) can be used as a proxy of strength-of-recommendation, such that:

– A therapy (or therapy combination) with higher LOE is more likely to have established evidence about efficacy than ones with lower LOE.
– A therapy (or therapy combination) with higher LOE is more likely to be accessible.
– A therapy (or therapy combination) with higher LOE from a drug that shares the same pharmacodynamic target (e.g., within the same drug class or have the same biological target) are more likely to lead to therapeutic significance.
- A therapy (or therapy combination) using the same biomarker, with efficacy in a different cancer types, is more likely to achieve a favourable therapeutic outcome.

Figure 1 illustrates a six-step process that constitutes the recommendation framework for inferring treatment options, derived using the task decomposition method.

**Figure 1.**
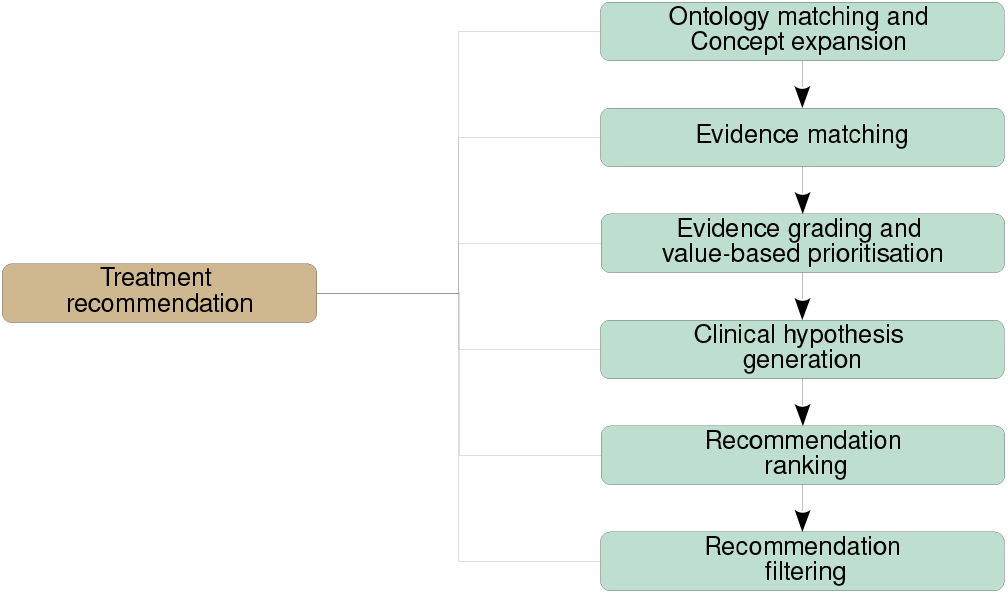
Proposed sequence of actions for inferring treatment options based on clinico-pathologic characteristics and biomarkers. (1) *Concept expansion by ontology matching*: The clinical and pathology concepts (including disease type and gene variant) are first expanded by using an established ontology or equivalent algorithm. (2) *Evidence matching*: The expanded ontology is matched to a knowledge base containing a pre-assessed of evidence with corresponding tier. (3) *Evidence grading and value-based prioritisation*: The level of evidence (LOE) of a matched entry is recorded by therapy (or therapy classes). Therapies are then ranked by a selected clinical value system. (4) *Clinical hypothesis generation*: the task of therapy recommendation is considered as formulating clinical hypotheses, prioritised based on a given value-based framework. For example, predicting the likely drug efficacy is essential to decide a potential list of therapies, providing an explainable basis for recommendation, and thus to identify the most valuable list of therapies that maximises theoretical benefit for the patient. (5) *Recommendation ranking*: given that multiple matched therapies to a clinicopathologic profile are often available, therapies should be rationally prioritised based on the same value framework stated above. (6) *Recommendation filtering*: manual filtering by physician is needed to remove non-relevant recommendations.

##### Structure of the IA

The IA is oriented towards multiple resolution closures that completely list *all* potential therapies. A non-learning IA is preferred in this setting (over a data-driven model), given the sparsity of efficacy data needed for inductive modelling. The “rule-of-thumb” models that are purely based on inference of biological pathways (and do not assess the established evidence) constitute the lowest strength-of-recommendation [18]. Recommendation based on established clinical data should be prioritised over *ab initio* rules, conforming to the accepted evidence-based decision paradigms.

##### The task and task environment

The objective of the IA is to sequentially optimise a predefined performance measure, with the most relevant one being the *strength-of-recommendation*. Within the boundary of curated knowledge, the *task environment* that the IA operates in, can be encompassed into a deterministic and fully observable close-world system comprising states and knowledge base rules. The *sensors* of the IA first transform clinical characteristics and biomarker profiles into propositional or continuous variables that form the initial state-space. The *goal* of decision inference, which form the IA’s *actuators*, is to produce a fully enumerated list of therapy options ranked by the associated strength-of-recommendation.

#### 3.1.2 Output requirements of an oncology therapy recommendation system

##### Evidence grading for producing prioritised recommendations

Drug recommendations should be prioritised based on the perceived clinical value of a treatment. To identify the best options in a treatment-refractory setting, the TRS should exhaustively list all suitable options that satisfy the patient’s clinicopathologic characteristics before prioritisation.

The IA should score each treatment recommendation using a predefined clinical value framework. For example, a novel but yet established therapy (e.g., drugs in Phase 1 clinical trials) should be deferred if a more established therapy option is available. In accordance with the goal of IA, recommendations are then ranked from most to least amount of evidence, to be followed by filtering by a physician.

##### Evidence synthesis: aggregated assessment of supporting literature

Multiple pieces of literature evidence need to be synthesised to determine its highest LOE to inform decisionmaking. A drug approved by regulatory authority (e.g., Food and Drug Administration, FDA) should be prioritised over investigational drugs. In general, positive results from late phase (e.g., phase 3) clinical trials should weigh more than early clinical trial data. Where there is a conflict of evidence, a summary of clinical values should be displayed to highlight the conflicting data.

##### Inference of efficacy and sensitivity of therapies

A decision should be made about whether a therapy is likely to benefit a patient, based on aggregated evidence about its sensitivity and resistance in the patient’s condition. The determination is also specific to the patient’s clinical history, such as prior therapies: a patient who has had exposure to a drug may be less likely to respond to another drug from the same class, of which should be deprioritised or filtered.

##### Modularisation of knowledge sources

The inference mechanism of TRS should be independent from, and capable of using alternative knowledge sources (such as ontologies and evidence knowledge base). To facilitate a modular design, the reasoning algorithm should be separated from content of the knowledge sources.

##### Transparency of inference process

The entire decision-making process should be fully explanatory to clinical stakeholders. Such transparency will ensure that peer-review of the intermediate steps can take place for explicit feedback. The TRS should inform user about each step of inference, for example, how pathogenicity is determined (e.g., variant calls as part of molecular pathology process) or how evidence is used in the reasoning process.

For options that do not have a sufficient evidence base for recommendation, the clinical hypothesis used for making the recommendations should also be explicitly stated by the TRS.

##### Customisable post-inference filtering

The therapy options inferred by the IA should be filtered for generating final outputs for clinical use. For example, a TRS should provide a mechanism to allow a physician user to remove options that are deemed ineffective or not accessible.

##### Interoperability

The recommendations produced by TRS should align with existing clinical practice guidelines. The outputs of a TRS should be integrated with a reporting system, and interoperable with specialised user interfaces, electronic medical record (EMR), and computerised physician order entry (CPOE) systems.

### 3.2 Knowledge representation and inference mechanism

Clinicopathologic characteristics and biomarkers can be readily encoded by a symbolic representation. The oncology domain knowledge can be represented by constraints defined by production rules in curated knowledge bases.

To avoid bias or omission in the resulting recommendations, the IA should find all consistent solutions in a *constraint-satisfying problem*. After the global search of solutions, the most valuable solutions are selected by a user-defined value function (e.g., strength-of-recommendation measure), before the post-inference filtering process.

#### Initial state space

The initial state space is described by a set of states in logical conjunction, based on clinicopathologic characteristics and biomarker profile. For example:

- Age: 65 years old
- Cancer type: Infiltrative ductal adenocarcinoma of breast
- Stage: IV
- Oestrogen receptor: positive
- *ERBB2*: amplified
- Prior therapy: Trastuzumab

#### Knowledge bases and rule-based inference

Symbolic reasoning is used to represent domain knowledge following the initiation of state space. The domain knowledge is explicitly encoded by IF-THEN production rules. For example:

- Sensitive to: MEK inhibitor → Sensitive to: Cobimetinib
- Oestrogen receptor: positive ∧ cancer type: Breast cancer → Sensitive to: Tamoxifen
- *ERBB2*: protein expression ∧ cancer type: gastroesophageal junction cancer → treatment: Cisplatin + Fluorouracil + Trastuzumab (LOE: 1)

The task of treatment recommendation is sequential in nature (e.g., determination of pathogenicity of a variant is needed before inferring sensitivity of drug). The reasoning process starts by matching the states to the left-hand-side (LHS) of rules.

A typical LHS may consist of combination of cancer type (e.g., “colorectal cancer”), clinical prerequisite (e.g., “no prior therapy”), pathology biomarkers (e.g., “mismatch repair deficient”), and specifies the state of biomarker (e.g., *“BRAF* V600E”), in logical conjunction. The initial states are incrementally expanded by matching the states against all *modus ponen* rules listed in knowledge bases. First-order logic can be used to expand the breadth of search, by inferring the relationship between states using curated ontologies of cancer types, biomarker classes, and drug classes.

A LHS propositional clause, which is expressed in conjunctive normal form, comprises antecedents in first-order patterns, whereas the logical successor function is defined by the right-hand-side (RHS) of a rule, which incrementally asserts new states when the LHS constraints are fully satisfied. Resolution closures are declared when no more new states can be expanded.

All solutions are reasoned using *exhaustive uninformed search*. This is preferred to avoid biased decision making: a complete set of goal states (therapy options) should be evaluated. Conversely, greedy, stochastic, heuristics (including A* algorithms) are not suitable for the purpose of inference as search space would be truncated.

#### Justification of forward-chaining approach

Reasoning by forward-chaining is preferred (over backward-chaining), as this inference mechanism gives a complete set of solutions. An additional advantage of forward-chaining is that it avoids repeated traversal of known states within a close-world system. Forward-chaining is also an efficient approach in this task, given that the search space is wide (i.e., many clinical scenarios, genetic variants, and cancer types) but relatively shallow with respect to the depth of inference (only few steps to reach the goals).

#### 3.2.1 Evidence aggregation for identifying therapy options

Several curated knowledge bases have been established to assess the quality-of-evidence about biomarker-driven therapies [19]. These knowledge bases, which categorise biomarkers and therapies with references to “actionability” (e.g., OncoKB [20], Precision Medicine Knowledge Base [21], Clinical Interpretation of Variants in Cancer [22], and Jackson Laboratory Clinical Knowledge Base [23]) have simplified the process of literature search. Different level-of-evidence (LOE) systems — scales that summarise drug maturity by formally assessing published literature — have been applied to facilitate knowledge synthesis with respect to significance of biomarkers [24,25] or therapies [26,27]. A recent development of a consensus therapy scale, European Society of Medical Oncology (ESMO) Scale of Clinical Actionability (ESCAT), has also been formed to reduce misinterpretation of gene sequencing reports, tumour board recommendations, and scientific communication [18].

##### Inference tracking

To assess the evidence quality associated with a recommendation, the IA should record how a recommendation is rationally derived along the path of inference when resolving a goal. To allow backtracking of ancestral states, all states in the LHS used for inferring the subsequent states should be recorded.

The proposed inference tracking mechanism provides a functionality to record the strengths-of-recommendation of states along the path of inference. To illustrate this concept, consider the following rule set:

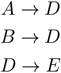

For the initial state *A* with strength-of-recommendation *V*(*A*), the subsequent inferred states can be recursively propagated using all ancestral states that satisfy the LHS of at least one *modus ponens* rule, such that:

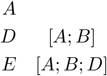

where […] are the recorded ancestral states from previous traversals. To determine the strength-of-recommendation based on the final state *E*, its LOE can be evaluated as a function of all antecedent states such that:

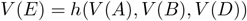

where *h*(*V*_1_(*t*), *V*_2_(*t*),…, *V_k_*(*t*)) is the heuristic function for determining the overall value of a therapy.

##### Evidence aggregation by merging strengths-of-recommendation

As discussed in Section 3.1.2, a simple yet useful heuristic in oncology practice is to select therapies with highest clinical values *V*(*t*), which form the strength of recommendation such that:

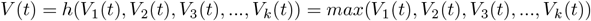

for 1 … *k* values associated with entries associated with a therapy or clinical trial *t*, and *h*(…) is the heuristic function. Using LOE as a proxy of strength-of-recommendation, the overall value of therapy *V* can be tracked along the inference path.

In the case that a therapy has ≥1 supporting piece of evidence, a summary measure represented by the best LOE can be calculated by maximising *V*(*t*) across all therapies:

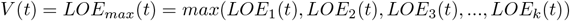

for 1 … *k* evidence entries for therapy or clinical trial *t*.

##### 3.2.2 Prioritisation of therapy options

Across the range of available drug options, the needs for prioritising molecular alterations to guide recommendations have been identified [26]. Given a set of therapies (and/or clinical trials options) **T** that eventually satisfies a set of clinicopathologic constraints, the options *T*_1_, *T*_2_,…, *T_n_* can be ranked according to clinical value function such that:

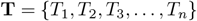

where *V*(*T*_1_) ≥ *V*(*T*_2_) ≥ *V*(*T*_3_) ≥ … ≥ *V*(*T_n_*).

This process guarantees the identification of a recommendation (or recommendations) with the highest clinical value, matching oncologist’s heuristics when selecting the most relevant therapy from a list of potential options such that:

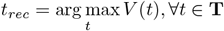

##### 3.2.3 Tracking information gaps during reasoning

###### Handling of incomplete information during the reasoning process

The above inference pipeline requires a completely deterministic scenario where all antecedent states are known prior to reasoning. However, therapeutic knowledge and real-world clinical data invariably suffer from a varying degree of unknowns, omission, and errors. Two scenarios are readily identified where incomplete data can be relevant to decision inference:

###### Incomplete assessment (unknown initial states)

Real-world medical data often contains many “missing values”, due to clinical constraints or limited information gathering. In these cases, potentially valuable options may not be fully identified by the TRS due to unmatched prerequisites.

###### Missing knowledge (evidence gap)

True “evidence” gaps and incomplete database coverage are well-known problems in precision oncology knowledge bases [28,29]. These gaps can also result in missed recommendations. For example, a targetable driver mutation that is yet to be curated but predicted to be oncogenic *in silico*, is otherwise unable to be matched to a therapy, due to lack of a curated production rule. Rationally inferred states can help bridge these gaps and allow useful recommendations to be made (Table 1).

**Table 1.** Examples of certain and inferred states using *ERBB2* alterations as an example

| Characteristics | Evidence-based (Certain) | Not evidence-based (Inferred) |
| --- | --- | --- |
| Oncogenicity | Well-characterised variants with determined oncogenicity (e.g., *ERBB2* L755S) | Variants inferred based on mutational position in the absence of strong clinical or pre-clinical data. (e.g., *ERBB2* L755A) |
| Gain-of-function mutation | Gene alterations where the mutation effects have been characterized. (e.g., *ERBB2* amplification) | Gene alterations where the likely mutation effects have been characterized, based on rules-of-thumb (e.g., *ERBB2* V777M) |
| Treatment | A specific treatment or treatment combination where Trastuzumab Emtansine (T-DM1) in the treatment of breast cancer | Trastuzumab Emtansine in non-breast solid tumours |
| Treatment class | Anti-HER2 antibody-drug conjugate in breast cancer | Any anti-HER2 antibody-drug conjugate in non-breast, non-gastric cancers |

###### Drug repurposing: rational extrapolation of recommendation

A rationally formed clinical hypothesis can be judiciously applied, which may then lead to valuable therapy options in patients who would otherwise have no other treatment options. Rational extrapolation of options forms the basis of *drug repurposing*. Without such a mechanism, the outputs of TRS would be strictly limited to the options concordant to the curated database, hindering its applicability.

###### Tracking extrapolated recommendations

As above, both “evidence-based” and “rationally inferred” recommendations are reportable. However, physicians inherently have a lesser degree of confidence in the latter type of recommendations. During the decisionmaking process, it is thus required for physicians to be informed if (and how much of) a particular recommendation is “evidence-based”. Computationally, there is a need to track which therapy options are based on extrapolation, to allow subsequent filtering and ranking.

The inference tracking method described in Section 3.2.1 can be used to label the hypothetically inferred states in this fully observable task environment. For instance, given a finite chain of inference below:

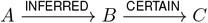

The rule *A → B* is triggered to allow intermediate state *B* to be asserted, which is then propagated to allow resolution closure at the terminal state *C*. To represent an inferred state, the history of all inferred path can also be labelled, such that:

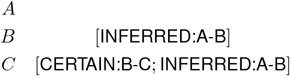

###### Summarising the extent of extrapolation in treatment recommendations

Extrapolation by assertion of missing states can take place at different steps of inference (e.g., hypothesising pathogenicity of a variant, repurposing a drug based on efficacy in another cancer type). To succinctly represent the information gaps, a simple summary measure of *level of matching* (LOM) is devised (Table 2). The LOM provides a semi-quantitatively way to summarise whether the terminal state is evidence-based or inferred from incomplete prior knowledge.

**Table 2.** Proposed definition of level-of-matching (LOM) to measure Examples of certain and inferred states using *ERBB2* alterations as an example

| Level of matching | Inference graph | Post-reasoning annotation of $C$ | Example |
| --- | --- | --- | --- |
| 1 - Complete match | $A \xrightarrow{\text{CERTAIN}} B \xrightarrow{\text{CERTAIN}} C$ | CERTAIN:A-B<br>CERTAIN:B-C | Using T-DM1 in HER2 amplified breast cancer |
| 2 - Partial match | $A \xrightarrow{\text{INFERRED}} B \xrightarrow{\text{CERTAIN}} C$ | INFERRED:A-B<br>CERTAIN:B-C | Using T-DM1 in HER2-amplified carcinoma of unknown primary |
| 3 - Completely inferred | $A \xrightarrow{\text{INFERRED}} B \xrightarrow{\text{INFERRED}} C$ | INFERRED:A-B<br>INFERRED:B-C | Using T-DM1 in carcinoma of unknown primary with *ERBB2* V777A mutation |

Level of matching (LOM) provides a summary measure of evidence gap that is used in inference. The LOM is designed to identify reasoned options that contain uncertainty during inference. A level of “complete match” (LOM 1) indicates that a therapy is fully inferred based on evidence, whereas “completely inferred” (LOM 3) indicates that the therapy is completely inferred without evidence base. In the case where the inferred therapy is partially based on evidence, a LOM 2 is designated.

#### 3.2.4 Computational complexity analysis

The proposed framework, using breadth-first strategy with forward-chaining, guarantees the completeness and optimality of the solution. It has the theoretical worst-case scenario of *O*^(*bd*+1)^ for both time and space complexities, where *b* is breadth, and *d* is the depth of the search.

### 3.3 Implementation of a treatment recommendation system in precision oncology

To illustrate the utility of this decision-support framework, a TRS was built for interpreting the results of tumour genomic profiling and clinicopathologic features (Figure 2) to provide decision-support in the tasks of: (1) ranking potential therapies based on a selected clinical value framework system, after considering all available evidence and interactions between genomic alterations; (2) predicting clinical drug sensitivity and resistance; and (3) preferential ranking of clinical trial options based on biomarker-matched evidence, using the same value tiering system above.

**Figure 2.**
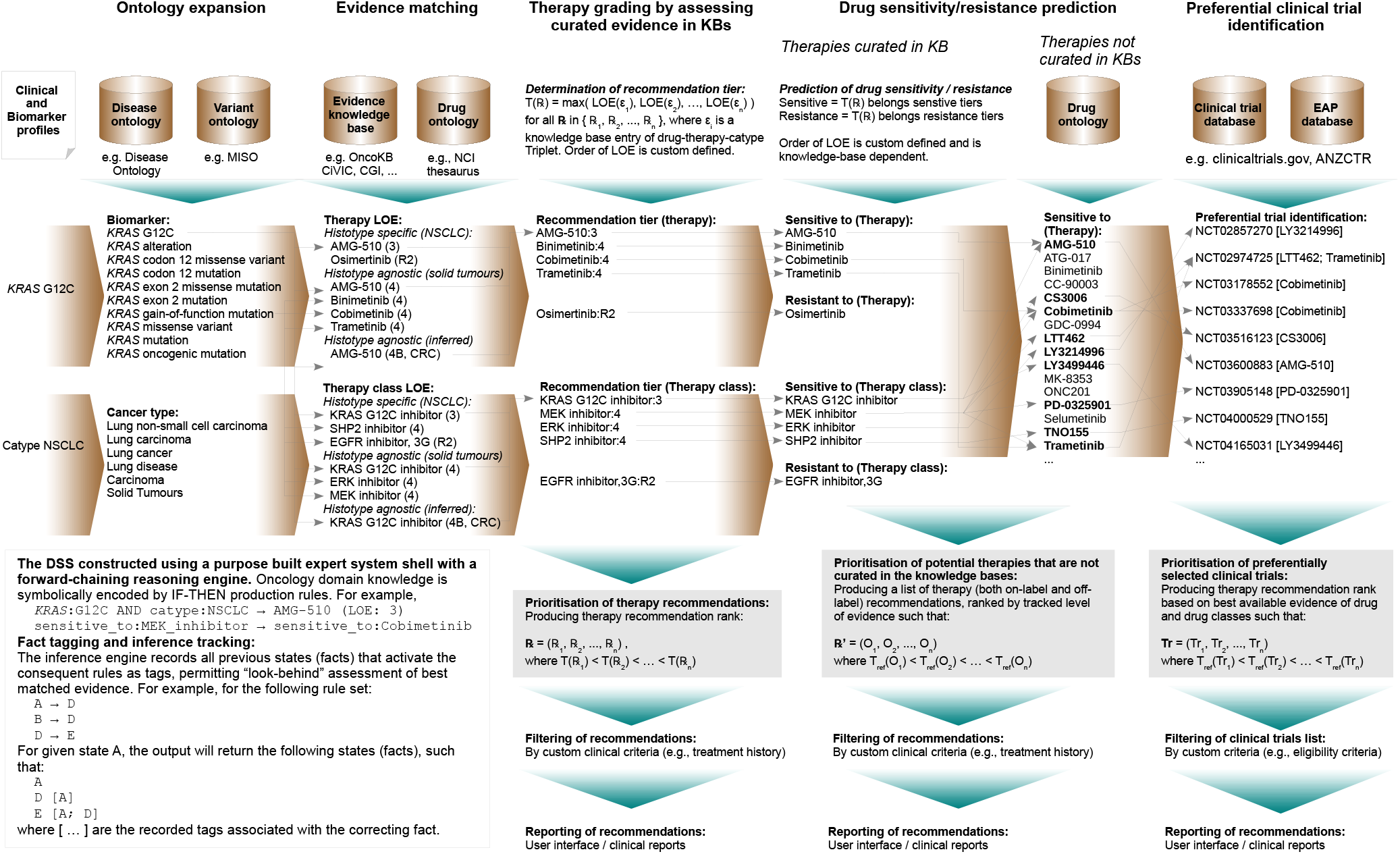
Overview of Precision Oncology Treatment and Trials Recommender (POTTR): a decision-support system that implements the proposed treatment recommendation framework. POTTR performs evidence-based inference of drug treatment and clinical trial options by making inference based on patient’s results of genomic biomarkers and clinical characteristics.

#### 3.3.1 Components of the implemented TRS

The TRS employs a modular design, implementing the action sequence described in Figure 1. In addition, the following specific requirements were also captured and implemented:

##### Concept expansion module

A concept expansion pipeline was designed to expand the breadth of initial states (i.e., clinical and biomarker profiles) to improve the subsequent matching process. It comprises the following subcomponents:

##### Disease ontology

Hierarchical taxonomies, including *Disease Ontology*, were used to infer cancer type hierarchy (e.g., HER2-positive breast cancer *is-a* breast cancer *is-a* solid tumour *is-a* neoplasm) [30].

##### Variant grouping

Mutations of commonly encountered variant types are denoted by HGVS Recommendations for the Description of Sequence Variants [31]. A custom pipeline was implemented to process gene alterations and annotate mutation into *variant groups* (if exist). The annotation module is implemented in place of a more sophisticated ontology, such as Sequence Ontology [32], where concepts are more general and fixed.

##### Expansion of positional gene features

A pipeline was implemented to expand positional gene features (e.g., exon numbers of mutation). Amino acid codons are referenced to the canonical transcripts listed in genome-build Ensembl database (GRCh38) [33]. Manual annotations of selected gene features are performed for key oncogenic driver genes (e.g., *KIT* “kinase domain” or *EGFR* “exon 19 deletion”).

The TRS did not implement a function for classifying pathogenicity of variants or determining the functional consequence of a mutation (e.g., gain-of-function mutation). These features are required to be determined upstream of the TRS, although a set of user-defined annotation rules may be used to accomplish this task.

##### Drug ontology module

A manually curated taxonomical hierarchy for drugs was implemented to link the drug names with respect to drug classes. Synonyms of selected drugs were extracted from National Cancer Institute (NCI) thesaurus [34].

##### Knowledge base integration module

To produce an evidence-based assessment, a module that integrates knowledge bases was implemented to curate biomarker-disease-therapy triplets linked with a LOE score. An empirical scoring function to determine clinical value of evidence entry is defined. For example, when OncoKB is employed [20], the preferred scoring for therapy selection based on the hierarchy of LOE is equivalent to LOE with default ranking of *R*1 > 1 > 2 > 3 > 4

##### Drug sensitivity prediction module

In addition to the curated list, a module was designed to expand the potential list of therapies through extrapolation of the sensitivity and resistance of a therapy (and its combination), based on drug class. Here the sensitivity of a disease-biomarker subgroup is hypothesized from known sensitivity of curated drug classes. The clinical value of a therapy option is then approximated by LOE as recorded by the inference tracking, and the strength-of-recommendation is assigned by identifying the maximum LOE associated with antecedent states during inference.

##### Recommendation prioritisation module

This module prioritises the list of therapy by the maximal score before returning to filtering by clinical criteria and user, implementing the concept described in Section 3.2.2. The LOM (Section 3.2.3) is calculated to indicate whether extrapolation of oncogenicity of a variant, therapeutic evidence about a drug or drug class, and cancer types have taken place.

##### Clinical trial filtering and prioritisation module

A “molecular-informed” approach was implemented to identify potential clinical trials of interest, similar to the concept outlined by Johnson et al [11]. In brief, this approach selects clinical trials that include an investigational therapy, of which the drug classes have shown benefit as informed by a disease or biomarker (e.g., an anti-HER2 monoclonal antibody in *ERBB2* amplified cancers). The trials selected by drug classes are subsequently prioritised by the tracked strength-of-recommendation given an investigational therapy, based on the curated knowledge. Finally, cancer type and genotype matching are performed to exclude ineligible clinical trials, based on eligibility criteria specified by trial sponsors.

#### 3.3.2 Case studies of prioritised therapy recommendations produced by the TRS

To illustrate how therapy options are rationally prioritised by the TRS, case studies of therapy prioritisation in three different cancer types with *ERBB2* alterations are shown in Table 3, using OncoKB as the knowledge base (Accessed May 2019) [20].

**Table 3.** Examples of rationally prioritised therapies inferred by POTTR

| Therapy | LOE | Matched evidence |  |  |  | LOM |
| --- | --- | --- | --- | --- | --- | --- |
|  |  | Oncogenicity | Drug | Drug class | Cancer type |  |
| **(1) HER2-amplified metastatic breast adenocarcinoma** |  |  |  |  |  |  |
| Pertuzumab + Trastuzumab | 1 | Matched | Matched | Matched | Matched | 1 |
| Trastuzumab | 1 | Matched | Matched | Matched | Matched | 1 |
| Trastuzumab Emtansine | 1 | Matched | Matched | Matched | Matched | 1 |
| Lapatinib + Trastuzumab | 1 | Matched | Matched | Matched | Matched | 1 |
| Neratinib | 1 | Matched | Matched | Matched | Matched | 1 |
| Lapatinib | 1 | Matched | Matched | Matched | Matched | 1 |
| Carboplatin + paclitaxel + trastuzumab | 3B | Matched | Matched | Matched | Inferred | 2 |
| DZD9008, Pyrotinib, ARX788, GQ1001, EO1001, Margetuximab, Tucatinib, Pertuzumab | - | Matched | Inferred | Matched | Inferred | 2 |
| **(2) HER2-immunohistochemistry negative metastatic breast adenocarcinoma but harbouring *ERBB2* V777L** |  |  |  |  |  |  |
| Neratinib | 3A | Matched | Matched | Matched | Matched | 1 |
| Trastuzumab Emtansine | 3B | Matched | Inferred | Inferred | Inferred | 2 |
| DZD9008, Poziotinib, Tucatinib, Pyrotinib, EO1001, Lapatinib, ARX788, GQ1001 | - | Matched | Inferred | Inferred | Inferred | 2 |
| **(3) *ERBB2* V777A in a carcinoma of unknown primary** |  |  |  |  |  |  |
| Trastuzumab Emtansine | 3B | Inferred | Matched | Matched | Inferred | 2 |
| Neratinib, Tucatinib, Lapatinib, Poziotinib, DZD9008, Pyrotinib | 3B | Inferred | Matched | Matched | Inferred | 2 |
| ARX788, EO1001, GQ1001 | - | Inferred | Inferred | Inferred | Inferred | 3 |

Abbreviations: LOE: Level of evidence (OncoKB); LOM: Level of matching.

In scenario 1 (HER2-amplified breast cancer), established therapies of standard of care were rationally prioritised as Trastuzumab with or without Pertuzumab, and Trastuzumab Emtansine, all of which were correctly identified as potential therapies. Investigational therapies were matched by matching known drug classes to new therapies, with specific therapeutic agents inferred, but deprioritised based on a lower LOE (e.g., pyrotinib and margetuximab).

In scenario 2 (a HER2-negative metastatic breast adenocarcinoma but found to harbour *ERBB2* V777L mutation), the “oncogenic mutation” grouping completely matched to Neratinib, whereas the option of “Trastuzumab Emtansine” was inferred and ranked lower.

In scenario 3 (*ERBB2* V777A in carcinoma of unknown primary) describes a scenario where oncogenicity of V777A is inferred from the amino acid position, and the sensitivity to anti-HER2 antibody-drug conjugate were inferred from other cancer types. In this scenario, the comparatively low LOE (i.e., untiered) indicates a lowest strength-of-recommendation. Overall, the LOM 3 is associated with investigational drugs of anti-HER2 monoclonal antibodies, implying the presence of evidence gaps.

## 4 Discussion

The main contribution of this paper is threefold. First, the decision framework presented here, together with the list of clinical premises, comprehensively models the oncologist’s decision-making process when selecting treatments for patients with advanced cancer. Second, the transparency of this informatic framework allows a consistent reasoning process in recommending treatments based on contemporary biomarkers. Third, the recommendation prioritisation process described here, which systematically applies knowledge of molecular biology using an evidence-based approach, synergises with the oncologist’s cognitive process in this task.

Further to the practical importance, this work also established a method of *inference tracking* in symbolic reasoning — which is a specialised form of cost search — that provides a transparent way to facilitate clinical evidence synthesis. In essence, this method provides the mechanism to rank and identify options with the highest clinical value for physicians. This method also allows assessment of quality of inference, which can be potentially applied to other decision-support tasks in another medical domain. Compared to other methods for representing uncertainty in reasoning (e.g., fuzzy logic), the proposed LOM score is also simple and should be readily understood by physicians.

More broadly, this framework illustrates how knowledge of cancer therapeutics described in the literature — central to evidence-based medical care — can be synthesised systematically with a symbolic reasoning system. Given the current state of precision oncology, using curated knowledge with propositional logic is preferred over data-driven methods: a primarily data-driven approach would not be feasible. There is a scarcity of patient-level data to allow inductive modelling of decisions (e.g., machine learning) in treatment-refractory cancers, notwithstanding the risk of selection and information bias. Therapies inferred only by real-world evidence are also considered hypothesis-generating only and cannot be considered as best clinical practice.

Several questions and limitations remain to be addressed, and these are subjects to active research. First, it remains to be determined how to best integrate an *in silico* intelligent frameworks to facilitate decision-making [13]. Second, an objective determination of prioritisation criteria in treatment recommendation is yet to be validated by formal comparative methods; semi-quantitative methods, such as an analytic hierarchy process, could potentially be employed for this task [35]. Third, the clinical benefits conferred by *in silico* TRS need to be elucidated. A thorough comparative evaluation with more biomarker-based treatment guidelines (e.g., the recent ESMO recommendation [6]) is desirable: it is hypothesised that a TRS can produce more specific drug recommendations than static guidelines, but the practical advantages about outcome remains unknown. Similarly, when combined with a comprehensive knowledge base, it is essential to study if an AI-based decision-support can provide more granular recommendations over tree-based algorithms or decision tables, which are predominantly used in traditional clinical trials [5]. Fourth, the intelligent framework may have a role in facilitating informatics-assisted clinical trial design to accelerate translational research. The role of TRS in clinical and translational research remains to be defined. Overall, formal clinical evaluations of this framework and associated tools, in both observational and implementation studies, will determine its benefit in patient outcomes.

## Software availability

The software implemented using this decision-support framework, Precision Oncology Treatment and Trial Recommender, is available as an open-source software on GitHub at https://github.com/fpylin/POTTR, under Apache Licence.

## Acknowledgement

The author declares no financial conflicts of interest. The author acknowledges the support from Prof. Richard Epstein and the Wolf family, without whom this work would not be possible. The author also thanks Dr. Subotheni Thavaneswaran, Dr. John Grady, and Prof. David Thomas for commenting on an implementation of POTTR.

